# Single-cell atlas of the first intra-mammalian developmental stage of the human parasite *Schistosoma mansoni*

**DOI:** 10.1101/754713

**Authors:** Carmen Lidia Diaz Soria, Jayhun Lee, Tracy Chong, Avril Coghlan, Alan Tracey, Matthew D Young, Tallulah Andrews, Christopher Hall, Bee Ling Ng, Kate Rawlinson, Stephen R. Doyle, Steven Leonard, Zhigang Lu, Hayley M Bennett, Gabriel Rinaldi, Phillip A. Newmark, Matthew Berriman

## Abstract

Over 250 million people suffer from schistosomiasis, a tropical disease caused by parasitic flatworms known as schistosomes. Humans become infected by free-swimming, water-borne larvae, which penetrate the skin. The earliest intra-mammalian stage, called the schistosomulum, undergoes a series of developmental transitions. These changes are critical for the parasite to adapt to its new environment as it navigates through host tissues to reach its niche, where it will grow to reproductive maturity. Unravelling the mechanisms that drive intra-mammalian development requires knowledge of the spatial organisation and transcriptional dynamics of different cell types that comprise the schistomulum body. To fill these important knowledge gaps, we performed single-cell RNA sequencing on two-day old schistosomula of *Schistosoma mansoni*. We identified likely gene expression profiles for muscle, nervous system, tegument, parenchymal/primordial gut cells, and stem cells. In addition, we validated cell markers for all these clusters by *in situ* hybridisation in schistosomula and adult parasites. Taken together, this study provides a comprehensive cell-type atlas for the early intra-mammalian stage of this devastating metazoan parasite.

## Introduction

Schistosomes are parasitic flatworms that cause schistosomiasis, a serious, disabling, and neglected tropical disease (NTD). More than 250 million people require treatment each year, particularly in Africa^1^. The life cycle of this metazoan parasite is complex. A schistosome egg hatches in water to release a free-living, invasive larva that develops into asexually replicating forms within aquatic snails (the intermediate host). From the snail, thousands of cercariae –a second free-living larval form– are released into freshwater to find and invade a mammal (the definitive host). In the mammalian host, the larvae (schistosomula) migrate and develop into distinctive male or female adult worms^2^ (Figure 1A). While the only drug currently available to treat schistosomiasis (praziquantel) works efficiently to kill adult parasites, it is less effective against immature parasites including schistosomula^3^. Understanding the parasite biology is a critically important step for developing novel strategies to treat and control this NTD.

**Figure 1.**
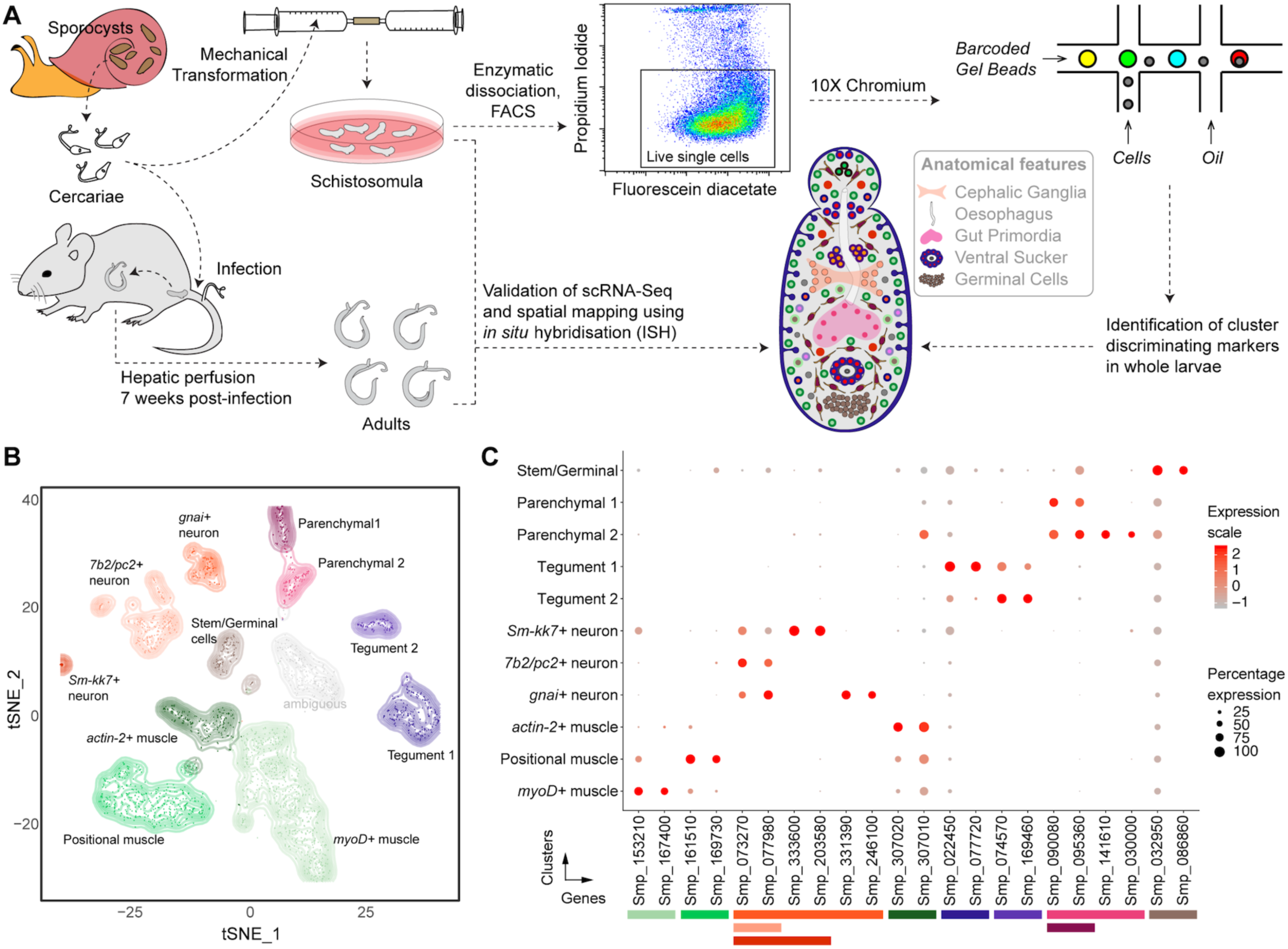
Identification of 11 transcriptionally distinct cell types in schistosomula. (A) Experimental scheme describing the sources of the parasite material, single-cell analysis and validation pipeline. Approximately 5,000 schistosomula per experiment were dissociated, followed by enrichment of fluorescein diacetate (FDA+) live cells using fluorescence-activated cell sorting (FACS). Cells were loaded according to the 1 OX Chromium single-cell 3’ protocol. Clustering was carried out to identify distinct populations and population-specific markers. Validation of population-specific markers was performed by in situ hybridisation (ISH). (B) t-distributed stochastic neighbour embedding (t-SNE) representation of 2,144 schistosomulum single cells. Clusters are coloured, distinctively labelled, and emphasised with density contours. One ambiguous cluster is de-emphasised and shown in grey. (C) Gene expression profiles of population markers identified for each of the cell clusters. The colours represent the level of expression from dark red (high expression) to light red (low expression). The sizes of the circles represent the percentages of cells in those clusters that expressed a specific gene. The colour bars under gene IDs represent the clusters in (B).

During invasion, the parasite undergoes a major physiological and morphological transformation from the free-living, highly motile cercariae to the adult parasitic form^2^. Upon penetration, the tail used for swimming is lost. Less than three hours after entering the host, the thick glycocalyx is removed and the tegument is remodelled to serve both nutrient-absorption and immune-protection roles^4^. Throughout the rest of the organism’s life in the definitive host, a population of sub-tegumental progenitor cells continuously replenish the tegument, allowing the parasite to survive for decades^5,6^. The schistosomula make their way into blood or lymphatic vessels and, one week after infection, reach the lung capillaries^7^. The migration through the lung requires coordinated neuromuscular activities, including cycles of muscle elongation and contraction^8^, to squeeze through capillaries and reach the general circulation^7^. Over the following weeks, the parasites mature further into sexually reproducing adults. Dramatic changes to the parasite are required that include posterior growth, remodelling of the musculature^9^ and nervous system^10,11^ as well as the development of the gonads^12^ and gut^13^. This extensive tissue development starts in the schistosomula, with stem cells driving these transitions^6,14,15^. However, to decipher cellular and molecular mechanisms underlying schistosomula development, a detailed understanding of the spatial organisation and transcriptional programs of individual cells are needed.

Important insights into major processes that underlie the transformations across the life cycle have been gained from bulk transcriptomic studies^5,6,14–24^. However, these studies are not able to quantify the relative abundance of different cell types from the absolute expression per cell, and the signal from highly expressed genes in a minority of cells can often be masked by a population averaging effect. Single-cell RNA sequencing has previously been used successfully to characterise cell types^25–32^ and understand how the cell expression profile changes during differentiation^31–38^. Notable examples include recent studies in the free-living planarian flatworm *Schmidtea mediterranea*^31,32,39^, a well-established model to study regeneration in the Phylum Platyhelminthes^40^.

Here, we have used scRNAseq to characterise two-day schistosomula obtained by *in vitro* transformation of cercariae^22^ using 10X Chromium technology and validated the cell clusters by RNA *in situ* hybridization (ISH) in schistosomula and adult worms. We identified eleven discrete cell populations, and described and validated novel marker genes for muscles, nervous system, tegument, parenchymal/gut primordia and stem cells. This study lays the foundation towards a greater understanding of cell types and tissue differentiation in the first intra-mammalian developmental stage of this NTD pathogen.

## Results

### Identification of 11 transcriptionally distinct cell types in schistosomula

We performed single-cell RNA sequencing of schistosomula collected two days after mechanically detaching the tail from free-living motile larvae (cercariae) (Figure 1A). To do so, we first developed a protocol to efficiently dissociate the parasites using a protease cocktail, after which individual live cells were collected using fluorescence-activated cell sorting (FACS) (Figure 1A and Supplementary Figure 1A). Using the droplet-based 10X Genomics Chromium platform, we generated transcriptome-sequencing data from a total of 3,513 larval cells, of which 2,144 passed strict quality-control filters, resulting in a median of 900 genes and depth of 283,000 reads per cell (Table S1). Given that an individual schistosomulum comprises ~900 cells (Supplementary Figure 1B), the number of quality-controlled cells theoretically represents >2x coverage of all cells in the organism at this developmental stage.

To create a cellular map of the *S. mansoni* schistosomula, we used a combination of the SC3^41^, Seurat^42^ and UMAP^43^ algorithms to cluster cells based on their mRNA expression levels and statistically identify marker genes that were best able to discriminate between the clusters (Figures 1B and 1C, Table S2-S5). To identify which cells each cluster represented, we curated gene set lists of previously defined cell-specific markers (Table S6). For example, tegument^5,6,44–46^ and stem^14,47–49^ cell clusters were identified based on known marker genes in *S. mansoni*, whereas muscle cells^50–52^ and neurons^53–55^ were identified based on characterised marker genes in mouse and humans (Table S6). Based on the marker genes identified using Seurat, we identified three distinct muscle-like clusters composed of 1,105 cells, two apparent tegumental clusters (253 cells), two parenchymal clusters (155 cells), one cluster resembling stem cells (94), three resembling the nervous system (311 cells), and one ambiguous cluster of 226 cells that could not be experimentally defined. Gene Ontology (GO) analysis of the marker genes generally matched the predicted cellular processes for each cluster (Supplementary Figure 1C). For instance, as expected, the stem/germinal cell cluster showed a significant enrichment in genes involved in translation, DNA replication, and RNA binding. Meanwhile, neuronal cells and muscle cells were enriched in processes involved in GPCR signalling and cytoskeleton, respectively. These analyses suggested that each cluster is molecularly distinct and likely display different biological functions. Therefore, we defined highly specific cluster-defining transcripts (potential cell markers) and characterised their spatial expression in both larval schistosomula and adult schistosomes by ISH (Table S7).

### Muscle cells show position dependent patterns of expression

Three discrete muscle clusters were identified by examining the expression of the well-described muscle-specific genes myosin^56^ and troponin^52^ (Figure 2A). One muscle cluster showed high expression of the uncharacterised gene Smp_161510, which was expressed along the dorso-ventral axis of two-day old schistosomula (Figure 2B). In adult worms, Smp_161510 exhibited no dorsal-ventral expression pattern; instead, Smp_161510 expression was scattered throughout the worm body (Supplementary Figures 2A and 2B). A subset of cells in this muscle cluster expressed *wnt* (Smp_167140) (Figure 2A). These *wnt*+ cells showed an anterior-posterior gradient in two-day schistosomula (Figure 2C) that remained consistent during the development from juveniles to mature adult worms (Figure 2D and Supplementary Figure 2A). Given that these markers have been shown to have distinct spatial distributions^57,58^, we decided to term this population ‘positional muscle’.

**Figure 2.**
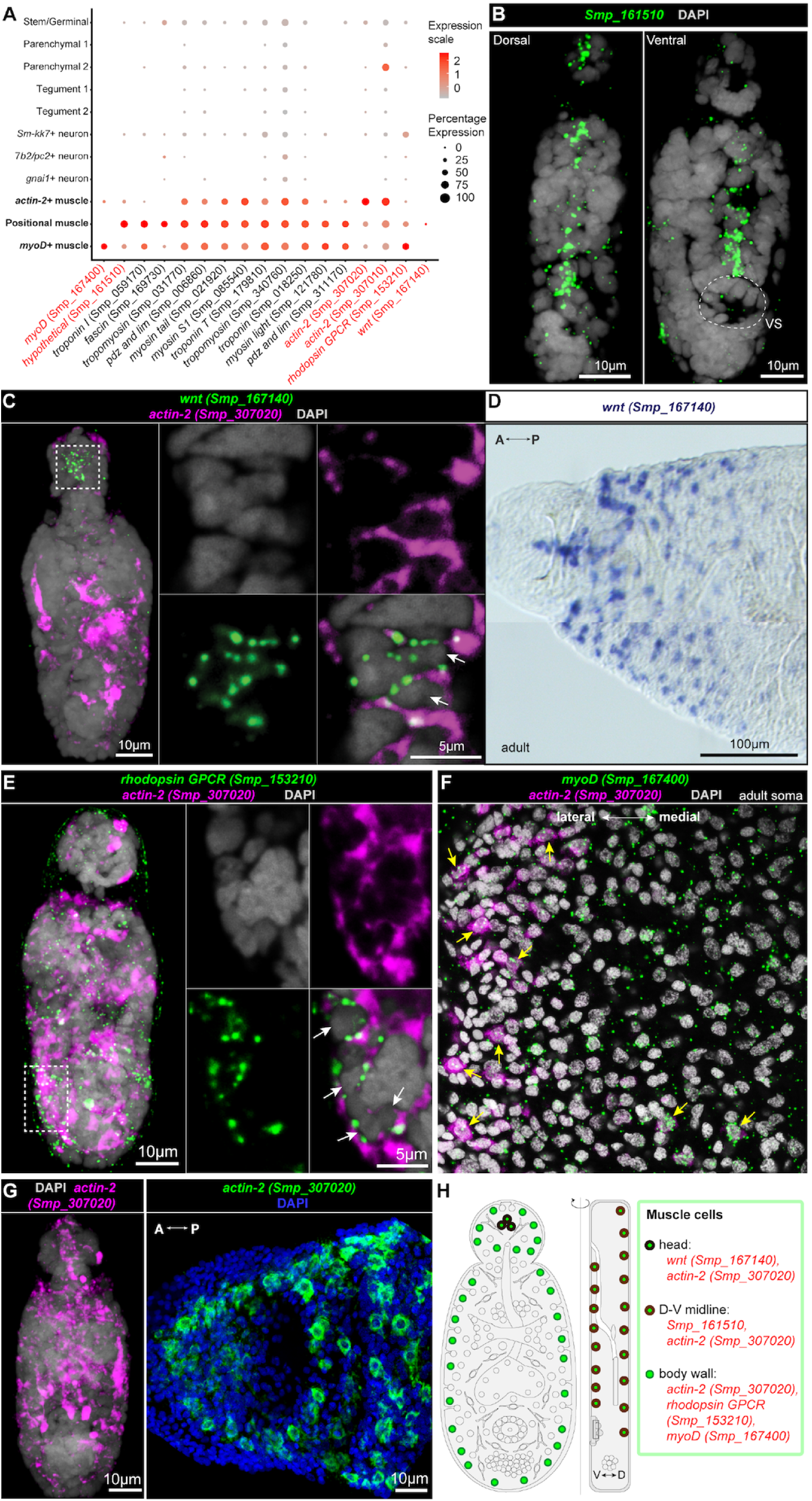
Muscle cells express positional information underlying parasite development. (A) Expression profiles of cell markers that are specific or enriched in the muscle clusters. Genes shown in red were validated by ISH. (B) FISH of Smp 1 61510. Smp 1 61510-expressing cells are found in dorsal and ventral sides along the midline. VS: ventral sucker. (C) Double FISH of *wnt* (Smp_l67140) and *actin-2* (Smp_307020). *wnt* is expressed in a subset of *actin-2+* cells in the head of the worm (white arrows). (D) Wholemount *in situ* hybridisation (WISH) of wnt in the head region of the adult worm. A: Anterior; P: Posterior. (E) Double FISH of *rhodopsin GPCR* (Smp_1 53210) with *actin-2* (Smp_307020). Left: MIP; Right: single magnified confocal sections of the dotted box. White arrows indicate doublepositive cells. (F) Double FISH of *myoD* (Smp_ 167400) and *actin-2* (Smp_307020) in adult soma. Scattered expression of *myoD* throughout the soma, with few double-positive cells (yellow arrows). (G) Spatial distribution of *actin-2* (Smp_307020) throughout the body of the parasite. Left panel: schistosomulum; Right panel: adult male. (H) Schematic that summarises the muscle cell types in 2-day schistosomula. Marker genes identified in the current study are indicated in red. All previously reported genes are shown in black. V: Ventral; D: Dorsal.

A second muscle-like cluster was uniquely found to express genes encoding a rhodopsin orphan GPCR (Smp_153210) (Figure 2E) and an ortholog of the myoD transcription factor (Smp_167400) from *S. mediterranea* (dd_Smed_v6_12634_0_1)^31^. Both genes showed a scattered expression pattern throughout the schistosomula (Figure 2E), and adult body (Figure 2F, Supplementary Figure 2B).

Finally, the third cluster of putative muscle cells was shown to be enriched in *actin-2* (Smp_307020, Smp_307010) expression. FISH confirmed *actin-2* expression throughout the body of the schistosomula (Figure 2A). Our transcriptomic data suggested that *actin-2* was enriched but not specific to this cluster. In line with the transcriptome evidence, ISH revealed that *actin-2* is also expressed in some cells of both the ‘positional muscle’ and *myoD*+ populations in schistosomula (Figures 2C, 2E-F) and in adults (Supplementary Figure 2C). Together, we identified three transcriptionally distinct cell types validated by ISH that represent schistosomula muscle cells (Figure 2H).

### Schistosomula have two distinct populations of tegumental cells

We identified two populations of tegumental cells (Tegument 1 and Tegument 2, Figure 3A). The first tegumental cluster (Tegument 1) expressed several known tegument genes, including four that distinguish it from Tegument 2 and encode: Fimbrin (Smp_037230), TAL10 (Smp_074460), Annexin B2 (Smp_077720) and Sm21.7 (Smp_086480) (Figure 3A; Table S6)^6,45,46,59^.

**Figure 3.**
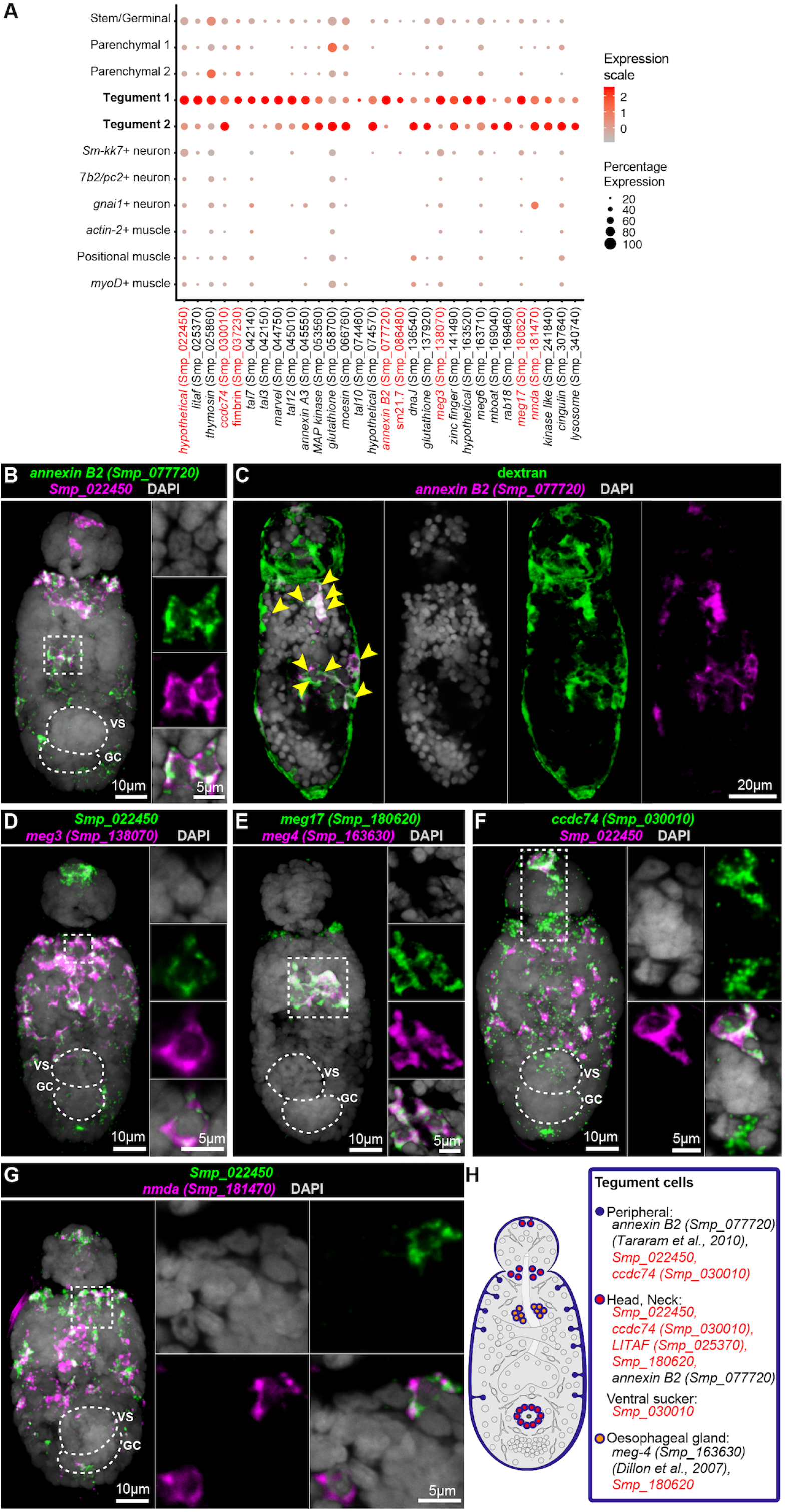
Two distinct populations of tegumental cells in schistosomula. (A) Expression profiles of cell marker genes that are specific of or enriched in the tegument clusters. Genes validated by ISH are marked in red. (B) Double FISH of Tegument I markers *annexin B2* (Smp_077720) and Smp_022450. The majority of the cells show co-localisation (white signal). MIP on the left, and zoomedin confocal sections on the right. (C) *annexin B2+* cells have taken up the fluorescent dextran. Yellow arrowheads indicate double positive cells. Single confocal sections are shown. (D) Double FISH of Smp 022450 and *meg3* (Smp_l 38070), both Tegument I markers. The majority of the cells show co-localisation (white signal). (E) Double FISH of *meg 17* (Smp 180620) with a known oesophageal gland gene *meg4* (Smp 1 63630). *meg 17* is expressed in other regions of the body including in the oesophageal gland. (F-G) Double FISH of Tegument 1 marker (Smp 022450) with (F) *ccdc74* (Smp_030010) and (G) *nmda* (Smp_l 81470). The majority of cells show co-localisation (white signal), while a subset of cells in the anterior portion of the worm show single positive cells for Tegument 2 markers. (H) Schematic that summarises the tegument cell populations in 2-day schistosomula. Marker genes identified in the current study are indicated in red. All previously reported genes arc shown in black.VS: ventral sucker; GC: germinal cell cluster.

The tegument 1 population also showed enrichment for an uncharacterised gene (Smp_022450) that, to our knowledge, has not previously been reported as a tegument-associated gene. We found that cells in the head, neck and body of the schistosomulum that expressed Smp_022450 co-localised with the tegument marker *annexin B2* (Smp_077720) (Figure 3B). In addition, cells expressing *annexin B2* and Smp_022450 were dextran+ (Figure 3C, Supplementary Figure 3A). Fluorescently conjugated dextran specifically labels tegumental cell bodies^6^, thus further validating Smp_022450 as a tegumental marker. In addition, Tegument 1 showed enrichment for microexon genes *meg3* (Smp_138070) and *meg17* (Smp_180620). The microexon gene *meg3* co-localised with the novel tegument gene Smp_022450 in the neck and anterior region of the larva (Figure 3D). The gene *meg17* was expressed in the neck and oesophageal region (Figure 3E). Given the expression of some *meg* genes in the oesophagus of adult male and female parasites^60^ and the developmental relationship between the oesophagus and the tegument^8,61^, we tested if *meg17* co-localised with any known oesophageal marker. We found that cells expressing *meg17* also expressed the known oesophageal marker *meg4*^62^ (Smp_163630) (Figure 3E). These results suggest that a subset of Tegument 1 cells likely represent primordial oesophageal gland cells.

Distinguishing the second tegumental cluster was challenging due a lack of Tegument 2-specific markers (Figure 3A). Two genes – *ccdc74* (Smp_030010) and *nmda* (Smp_181470) *–* with similarly enriched expression in both clusters were selected for further investigation. Double FISH experiments using either *ccdc74* (Smp_030010) or *nmda* (Smp_181470) with Smp_022450 showed co-localisation of expression (Figure 3F-G). In addition, these cells were also dextran+^6^, confirming their tegumental assignment (Supplementary Figure 3B and 3C). In adults, marker genes of Tegument 1 and 2 showed overall similar enriched expression patterns in the anterior cell mass, ventral sucker, and tegumental cells throughout the worm body (Supplementary Figures 3D and 3E).

To explore more subtle differences in expression profiles between these two tegumental populations, we investigated tentative functional differences. Analysis of marker genes for Tegument 2 using the STRING database predicted a group of interacting genes involved in clathrin-mediated endocytosis^63^ (Supplementary Figure 3F-3H). This group included genes that encode Phosphatidylinositol-binding clathrin assembly protein (Smp_152550), Epsin15-related (Smp_171640) and Epsin4 (Smp_140330) proteins. The potential involvement of Tegument 2 cells in calcium binding (Supplementary Figure 1C) and clathrin-mediated endocytosis is consistent with previous studies showing that numerous vesicles are produced by endocytosis from cell bodies and trafficked to the syncytial cytoplasm of the tegument^64,65^. Together, the evidence provided here supports these cells being part of the schistosomulum tegument (Figure 3H).

### Identification of schistosome parenchymal and primordial gut cells

Schistosomes, like other platyhelminths, are acoelomates and lack a fluid-filled body cavity. Instead, their tissues are bound together by cells and extra-cellular matrix of the parenchyma^20^. We identified two cell types that most likely represent parenchymal cells (Parenchymal 1 and 2) that showed enriched expression of numerous enzymes such as lysosome, peptidase, and cathepsin (Figure 4A).

**Figure 4.**
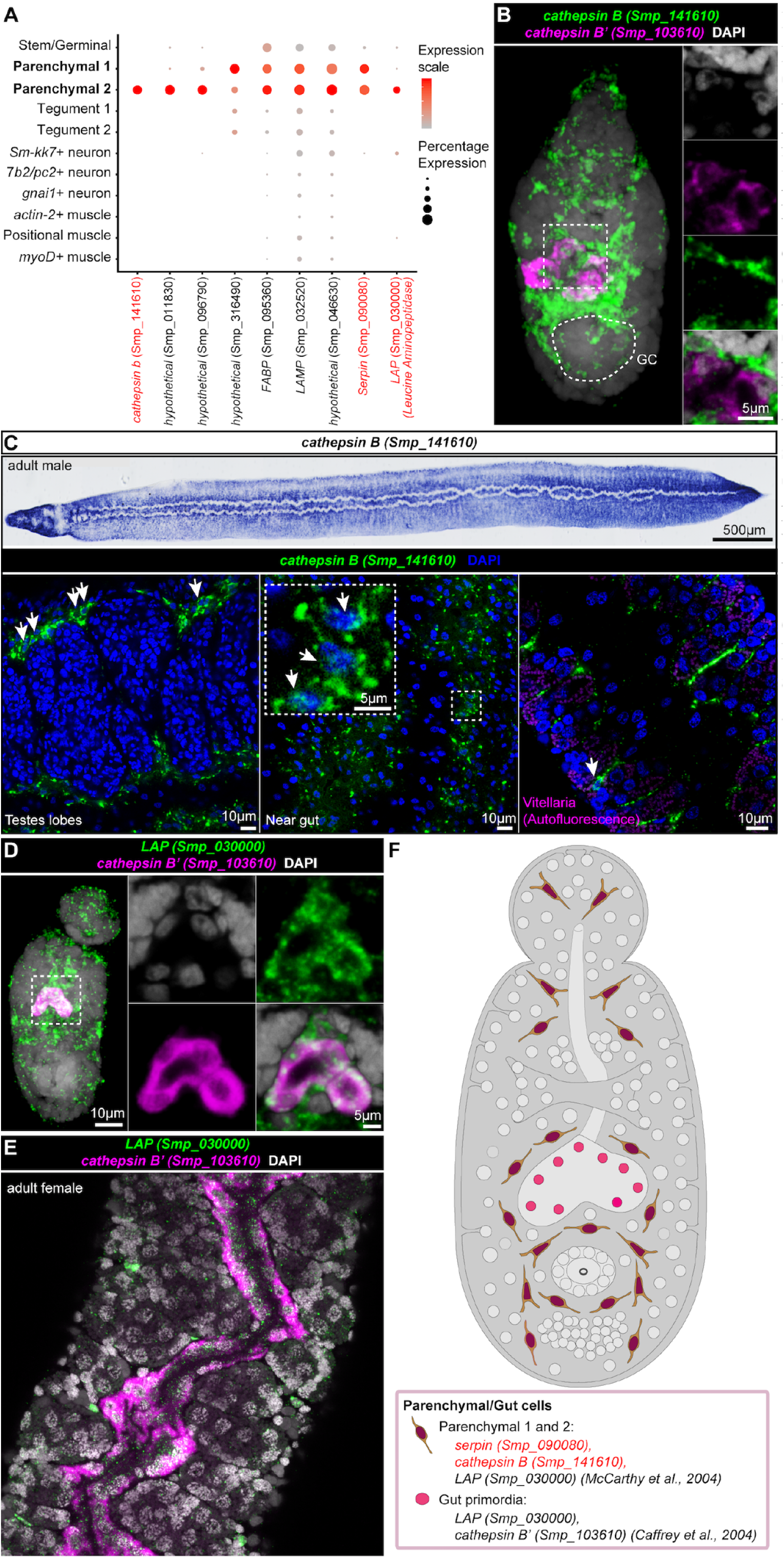
Identification of schistosome parenchymal and primordial gut cells. (A) Expression profiles of cell marker genes that are specific or enriched in the parenchymal clusters. Genes validated by ISH are marked in red. (B) Double FISH of parenchymal *cathepsin B* (Smp_l41610) with a known marker of differentiated gut, *cathepsin* B’(Smp_103610). No expression of parenchymal *cathepsin* Bis observed in the primordial gut. GC: germinal cell cluster. (C) WISH (top) and FISH (bottom) of parenchymal *cathepsin* Bin adult males. White arrowheads indicate positive cells in the bottom part of the figure. Single confocal sections shown for FISH. (D-E) *lap* (Smp 03000) is expressed in both parenchyma and in the (D) gut primordia as well as (E) adult gut, shown by double FISH with the gut *cathepsin B* (Smp_103610). (F) Schematic that summarises the parenchymal cell populations in 2-day schistosomula. Marker genes identified in the current study are indicated in red. All previously reported genes are shown in black.

Cells expressing *cathepsin B* (Smp_141610) were spread throughout the worm parenchyma and showed long cytoplasmic processes stretching from each cell (Figure 4B-C and Supplementary Figure 4D-E). A similar expression profile was observed for *serpin* (Smp_090080) expressing cells in the later stages of schistosomula as well as in adult parasites (Supplementary Figures 4A-4C). In addition, parenchymal cells did not co-express other cell type markers except for *actin-2*, which showed slightly overlap in expression (Supplementary 4F, 4J).

In Parenchymal 2 cells, we found that *leucine aminopeptidase* (*lap*) (Smp_030000) was expressed in the primordial gut (*cathepsin B*’(+) and surrounding parenchymal tissue (Figure 4D). Such mixed gut/parenchymal expression was also observed in adult parasites (Figure 4E, Supplementary Figure 4B). This is consistent with previous studies in adult parasites where LAP was detected in the gut and in cells surrounding the gut^66^. Overall, the identified genes that mark schistosomula parenchyma, while a few of them are also expressed in the gut primordia (Figure 4F).

### Stem cells in two day old schistosomula

Recently, it was shown that schistosomula carry two types of stem cell populations: somatic stem cells and germinal cells^15^. The somatic stem cells are involved in somatic tissue differentiation and homeostasis during the parasite intra-mammalian development, whereas the germinal cells are presumed to give rise to germ cells (sperm and oocytes) in adult parasites^15^. Less than 24 hours after the cercaria enters the mammalian host to become schistosomulum, ~5 somatic stem cells at distinct locations begin to proliferate^15^ (Figure 5B). Germinal cells, on the other hand, are thought to be packaged in a distinct anatomical location called the germinal cell cluster, and only begin to proliferate ~ 1 week after penetrating the host^15^.

**Figure 5.**
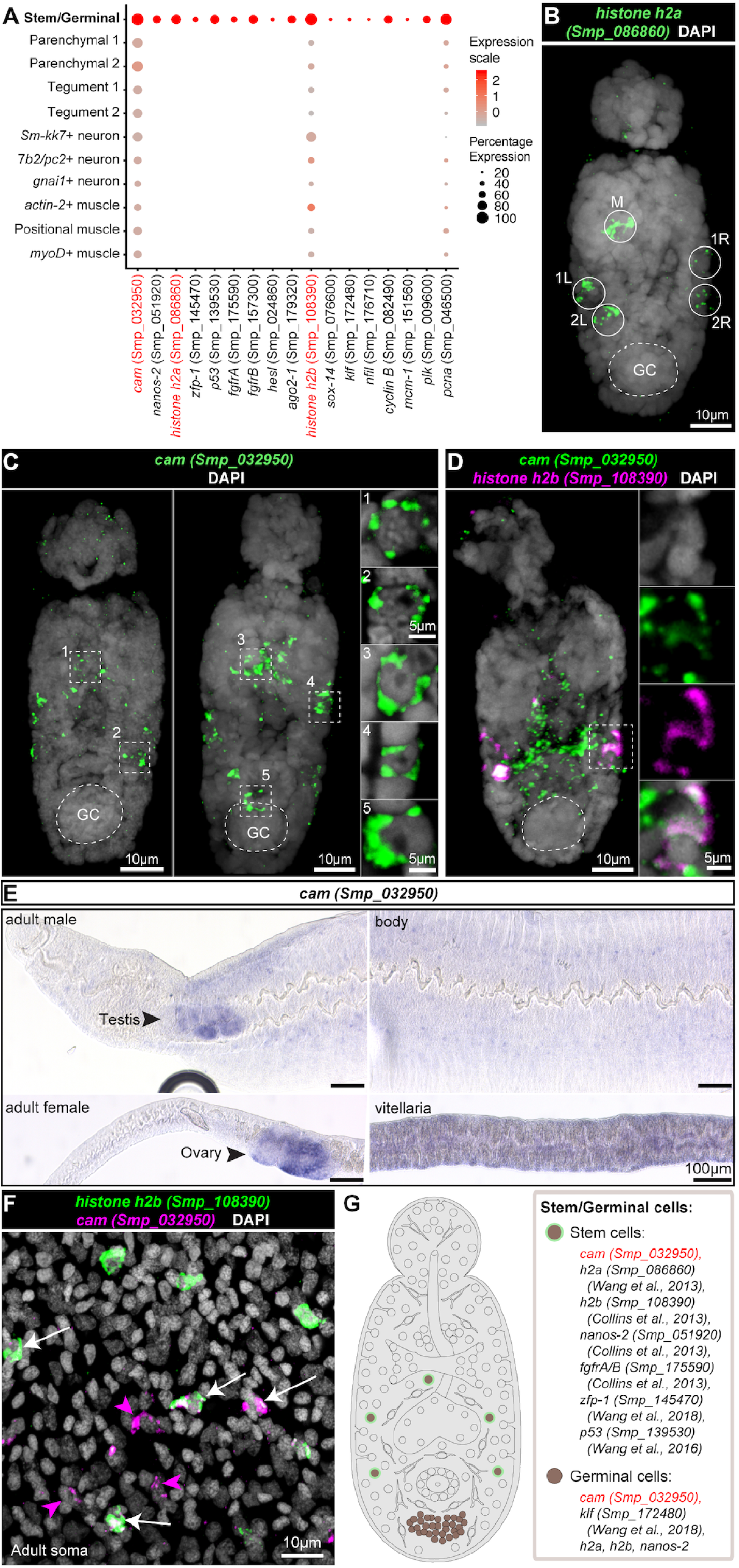
A single cluster of stem cells in 2-day old schistosomula. (A) Expression profiles of cell marker genes that are specific or emiched in the stem/ germinal cell cluster. Genes validated by JSH are marked in red. (B) FISH of *h2a* (Smp_086860) shows -5 stem cells located at distinct locations - 1 medial cell (M) and 2 lateral cells on each side (1 Land 2L, lR and 2R; L: left; R: right). (C) FISH of *calmodulin* (Smp_032950) shows a similar localisation pattern as *h2a*, with some worms with a few more *cam* I cells in the medial region as well as in the germinal cell cluster region. GC: germinal cluster (D) Double FISH of *calmodulin* (Smp_032950) and a previously validated schistosome stem cell marker *h2b* (Smp I 08390). (B-D) MlP is shown for the whole worms, and magnified single confocal sections are shown for the dotted box area. (E) WISH of *calmodulin* (Smp 032950) in adult parasites shows enriched expression in the gonads including testis, ovary, and vitellarium, as well as in the mid-animal body region. (F) Double **FISH** of *calmodulin* and *h2b* in adult soma. A single confocal section is shown. White arrows indicate co-localisation of two genes and magenta arrows indicate expression of only one gene. (G) Schematic that summarises the stem and germinal cell populations in 2-day schistosomula. Marker genes identified in the current study are indicated in red. All previously reported genes are shown in black.

We identified a single stem/germinal cell cluster that expressed the canonical cell cycle markers *histone h2a* (Smp_086860)^15^ and *histone h2b* (Smp_108390)^6^ (Figure 5A). In addition, this cluster also had a significant enrichment of translational components (Supplementary Figure 1C). We confirmed that *histone h2a* (Smp_086860) is expressed in ~5 cells, 1 medial and 2 sets on each side (Figure 5B) and also in the germinal cell cluster a few days later (Supplementary Figure 5A). In adults, *histone h2a* (Smp_086860) is expressed in somatic cells as well as in cells of the gonads (testis, ovary, and vitellaria) (Supplementary Figure 5B). In addition, we identified a novel stem/germ cell marker *calmodulin (cam)* (Smp_032950). This gene was expressed similarly to *h2a*, but in some schistosomula, a few more *cam*+ cells could be observed medially as well as near the germinal cell cluster (Figure 5C). The *cam*+ cells were also positive for *h2b* (Figure 5D), and found to be expressed in the adult gonads (Figure 5E) and in adult soma (Figure 5F).

In addition to *histone h2a* (Smp_086860), *histone h2b* (Smp_108390) and *cam* (Smp_032950), cells in this cluster expressed stem cell markers including *fgfrA* (Smp_175590) and *nanos-2* (Smp_051920)^14,15,67^ (Figure 5A). Given that many of these genes have been associated with two distinct stem cell populations^15^ (somatic and germinal), we tested if these cells could be further subclustered, but were unable to do so, presumably due to the low expression level of some of these genes in most cells in this cluster (Supplementary Figure 5C). Overall, these data suggest that this cluster does indeed represent population(s) of stem cells that might give rise to somatic and germ cells during the course of parasite development within the mammalian host (Figure 5G).

### Heterogeneity in cells of the schistosomulum nervous system

Platyhelminths have a central nervous system comprised of cephalic ganglia and main nerve cords, and a peripheral nervous system with minor nerve cords and plexuses^10^. This system also plays a neuroendocrine role by releasing neuromodulators during development and growth^10,68,69,70^.

We identified three distinct populations that expressed neural-associated genes (Figure 6A). One population was characterised by the expression of genes encoding neuroendocrine protein 7B2 (*7b2*, Smp_073270) and neuroendocrine convertase 2 (*pc2*, Smp_077980) and lack of *gnai* (Smp_246100) expression (Figure 6A). The *in situ* hybridisation of *7b2* (Smp_073270) showed expression in cells of the cephalic ganglia in schistosomula (Figure 6B-C). The cephalic ganglia region was identified using lectin succinylated Wheat Germ Agglutinin (sWGA)^11^ staining. In adult worms, *7b2* was expressed in the cephalic ganglia as well as in the main and minor nerve cords (Figure 6C). We refer to this cluster as ‘*7b2/pc2+* nerve’ cells.

**Figure 6.**
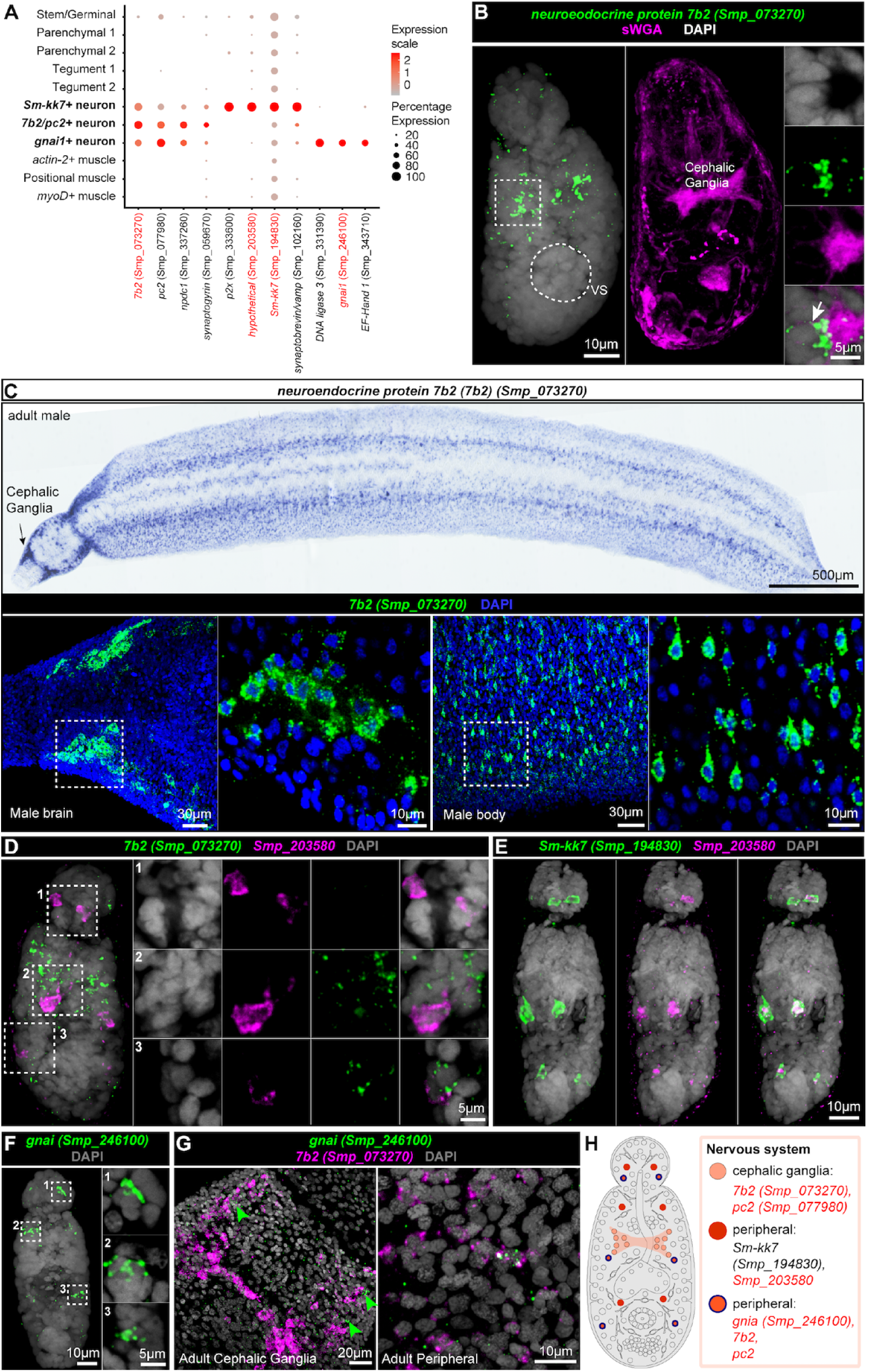
Heterogeneity in cells of schistosomula nervous system. (A) Expression profiles of cell marker genes that are specific or enriched in the neuronal clusters. Genes validated by ISH are marked in red. (B) Cephalic ganglia marked by sWGA lectin shows co-localisation with *7b2* (Smp_073270). White arrows indicate co-localisation of gene-sWGA lectin (C) WISH (top) and FISH (bottom) of *7b2* (Smp_073270) in adults. Single confocal sections are I shown for FISH. (D) Double FISH of 7b2 (Smp 073270) and Smp 203580 shows that six cells that are Smp_203580+ (in magenta) do not co-localise with *7b2* 1 cells (in green). (E) Double FISH of Smp_203580 with *Sm-kk7* (Smp_l94830). All Smp_203580+ cells co-localise with *Sm-kk7*. (F) *gnai* (Smp 246100) FISH shows expression in a few cells in the head and in the body region. (G) Double FISH of *gnai* with *7b2* shows co-localisation in the peripheral neurons. Single confocal sections are shown. (H) Schematic that summarises the neuronal cell populations in two-day schistosomula. Marker genes identified in the current study are indicated in red. All previously reported genes are shown in black.

A second population expressed the uncharacterised gene Smp_203580 (Figure 6D). Co-localisation experiments with *7b2* confirmed that this population was distinct from the central ganglia population (Figure 6D). In the larvae, only six cells (two cells in the head and four cells in the body) expressed the novel marker Smp_203580 (Figure 6D) but in adults, an expanded number of cells were found throughout the body of the parasite (Supplementary Figures 6A and 6C). These cells displayed 2-3 long cellular processes, branching into different directions (Supplementary Figure 6B). Interestingly, cells in this cluster also expressed the marker gene encoding KK7 (Smp_194830), known to be associated with the peripheral nervous system in *S. mansoni*^55^ (Figure 6E, and Supplementary Figures 6A and 6D). Therefore, we refer to this population as ‘*Sm-kk7+* nerve cells’.

Finally, we identified a population of cells that expressed *gnai* (Smp_246100), a gene encoding a G-protein G(i) alpha protein. FISH experiments showed expression of this gene in three cells: one in gland region of the head, one in the neck region, and one in the body region (Figure 6F). In adults, this gene is expressed around the main and minor nerve cords (Figure 6G and Supplementary Figure 6E and 6F). Some *gnai*+ cells are also *7b2*+ (Figure 6G). We designated this population as ‘*gnai+* neurons’. Overall, neuronal cells are transcriptionally and spatially heterogeneous (Figure 6H) and thus are expected to be involved in diverse biological processes.

### Conserved gene expression patterns in stem cells and neurons between *S. mansoni* and *Schmidtea mediterranea*

Given that some of the populations described herein had not been previously characterised, we asked if we could further annotate our dataset by comparison to previously annotated single-cell RNAseq data from *Schmidtea mediterranea*, the closest free-living model organism to *S. mansoni*^31^. To compare clusters, we used a random forest (RF) model trained on *S. mediterranea* to map gene expression signatures between both datasets^71^. Using the RF model, we classified each of the larval *S. mansoni* cells using the adult *Schmidtea* labels. We discovered that the stem cell population in our dataset mapped to *Schmidtea* stem cells (Figure 7). This is consistent with previous work that showed similarities between *Schmidtea* and *S. mansoni* stem cells^5,14,15,49,67,72^. We found that *Sm-kk7*+ cells in schistosomula mapped to the neuronal population annotated as *otoferlin 1* (*otf1*+) cells described by Plass *et. al*^31^. In addition, *7b2/pc2*+ cells in *S. mansoni* mapped to *spp11*+ and Chat neurons as well as neural progenitors in *Schmidtea* (Figure 7). In addition, tegument clusters in *S. mansoni* mapped to early and late epidermal progenitors in *Schmidtea*. The rest of the clusters in *S. mansoni* were labelled as *psd+* cells (of unknown function in *S. mediterranea*) and neoblasts. Taken together, these results suggest that despite great differences in developmental stages between larval schistosomula and the asexual adult *Schmidtea mediterranea* used for this comparison, marker genes for stem cells and neuronal populations have been conserved (Table S8-S10).

**Figure 7.**
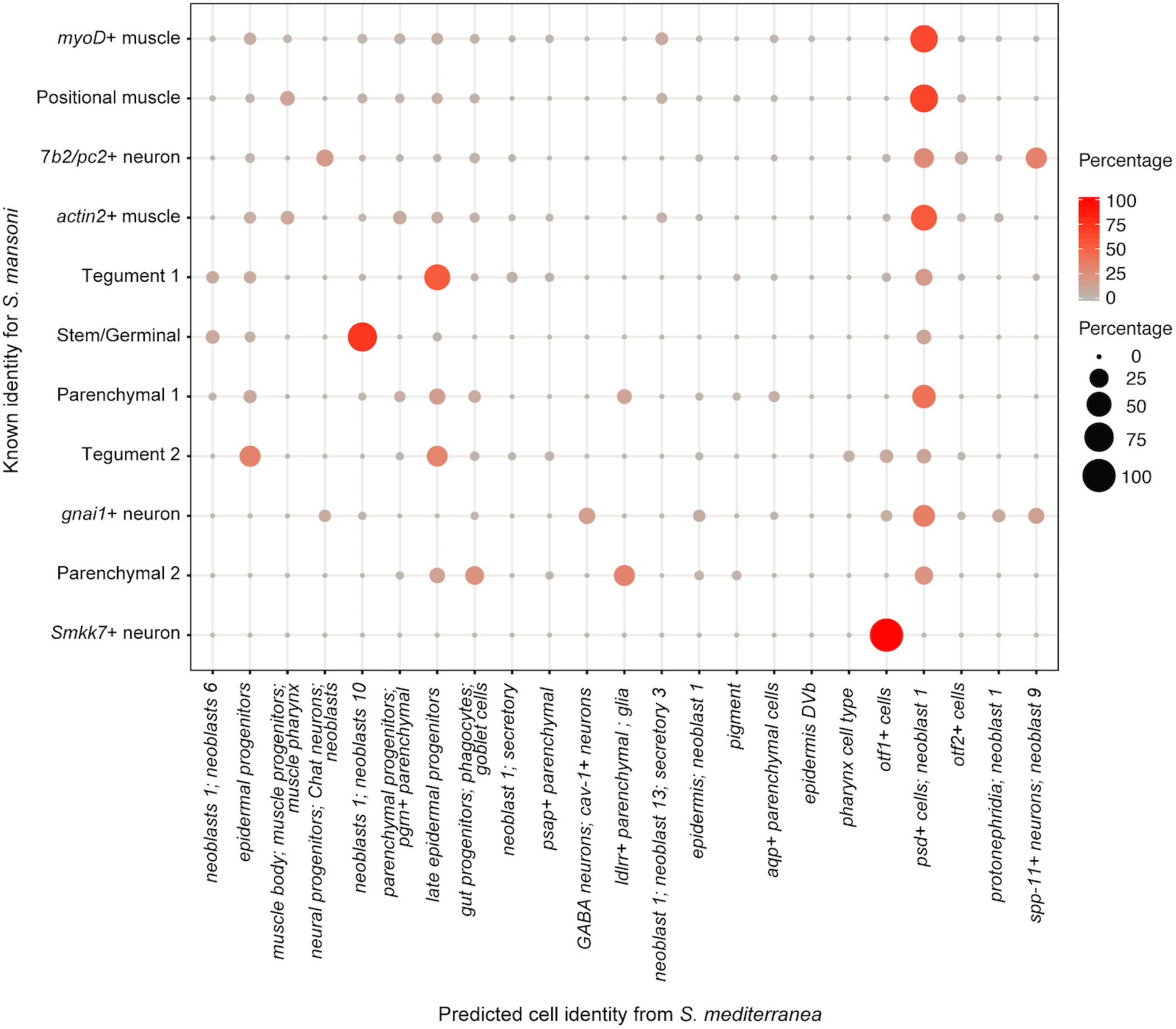
Conserve gene expression patterns in stem cells and neurons between *S. mansoni* and *Schmidtea mediterranea*. Dot plot showing the percentage of cells within each of the schistosomulum clusters (rows) that were mapped to *Schmidtea mediterranea* scRNA seq dataset **J1** (columns) using a multiclass random forest classifier (RF). The colours and size of the circles represent the proportion of cells assigned to a particular label. Large circles in red represent 100% of cells. The small circles in light red represent 0% of cells.

## Discussion

In this study, we have generated a cell atlas of the schistosomulum, the first intra-mammalian developmental stage of *S. mansoni* and a key target for drug and vaccine development^73,74^. Our transcriptome analysis enabled the characterisation of 11 distinct clusters, with sufficient sensitivity to detect as few as three cells per parasite, as demonstrated by the ISH experiments (Figure 6F). Importantly, the latter allowed us to validate key marker genes for each of the cell clusters, spatially mapping the cell populations in both schistosomula and adult worms and linking transcriptomic profiles to anatomical features of the organism.

By determining the transcriptome of individual cells from schistosomula, we uncovered marker genes not only for known populations, such as stem and tegument cells, but also for previously undescribed cell clusters, such as parenchymal cells. We found that marker genes of the parenchymal tissue are also expressed in the primordial gut. However, the relationship between the parenchyma and gut primordial cells is yet to be determined. In planarians, the orthologous *cathepsin* gene (dd_Smed_v6_81_0_1) is a marker for cathepsin+ cells that include cells in the parenchyma^32^. This planarian *cathepsin* (dd_Smed_v6_81_0_1) is also expressed in the intestine^32^ and gut phagocytes^31^. Similarly, planarian *aminopeptidase* (dd_Smed_v6_181_0_1) is expressed in *cathepsin*+ cells, epithelia and intestine^31,32^. Thus, further work is required to characterise schistosome parenchymal cells and their signaling mechanisms with the surrounding gut cells^75^.

Until now, *S. mansoni* cell types have been revealed primarily through a combination of morphological and ISH studies of specific tissues, with stem and tegument cell populations being among the best characterised^5,6,14,15,49^. In the present study, we identified and validated a novel stem cell marker *calmodulin* (Smp_032950) that, to our knowledge, has not previously been associated with stem cells. Calmodulins are Ca^2+^ transporters required for the miracidium-to-sporocyst transition, sporocyst growth^76^ and egg hatching^77^. In addition, we found this gene to be expressed in the reproductive organs of adult males and females.

Coordinated neuromuscular activity is essential for schistosomes to migrate through host tissue^78^. Although circular and longitudinal muscle layers have been described in *S. mansoni*^9,11,78^, we found no evidence that the three muscle clusters correspond to different anatomical fiber arrangements. In the free-living planarian *S. mediterranea*, a population of muscle cells also shows no specific muscle layer localisation, but instead forms a cluster based on enriched expression of position-control genes (PCGs)^32,79^. We therefore reasoned that this may be the case for at least some of the muscle cells in our dataset.

Knowledge of planarian stem cells has previously informed the study of stem cells in *S. mansoni*^67^. Our comparison between schistosomula and *S. mediterranea* clusters uncovered conserved features for stem cells and neurons and served to support cell type assignment in schistosomula. Given that nerve cell populations have remained poorly characterised at the transcriptome level in schistosomes, planarians may serve as a model to understand the nervous system biology in schistosomula. A particularly attractive feature of the planaria biology is the remarkable regenerative properties of the these worms. An individual worm comprises all cell types at intermediate stages of development and regeneration^31,32,80^. This has enabled recent single-cell sequencing studies in planarians to characterise developmental trajectories from within the soma of adult worms^31^. However, schistosomes do not share this regenerative property with their distant free-living relatives, instead intermediate stages of schistosome development necessarily need to be captured. The data from the present study represent the first logical step in that characterisation.

Despite having successfully characterised several previously unknown marker genes and populations, we faced challenges throughout the course of this study. Some cells were not detected, possibly because they are difficult to isolate or relatively rare. One notable example was the absence of eight known protonephridia cells in the parasite at this developmental stage^11,81^. Previous single-cell studies in *S. mediterranea* have found that relatively rare cell types are sometimes embedded in larger neuronal clusters^31,32^, and therefore, it is possible that this is also the case for this cell group. In addition, schistosomula obtained for this study were a mixture of males and females. While the male and female schistosomula are morphologically identical, they may bear transcriptomic differences that are important for early stages of reproductive development^82^. Future scRNA-Seq studies obtained separately from male and female schistosomula will be needed to resolve this question.

Our study demonstrates the power of single-cell sequencing, coupled with ISH validation, to transcriptionally and spatially characterise cell types of an entire metazoan parasite for the first time. This approach is essential for unravelling the developmental biology of this important parasite.

## Materials and methods

### Ethics statement

The complete life cycle of *Schistosoma mansoni* (NMRI strain) is maintained at the Wellcome Sanger Institute (WSI). The mouse infections at WSI were conducted under Home Office Project Licence No. P77E8A062 held by GR, and all protocols were presented and approved by the Animal Welfare and Ethical Review Body (AWERB) of the WSI, and Institutional Animal Care and Use Committees (IACUC) at the University of Wisconsin-Madison (protocol M005569). The AWERB is constituted as required by the UK Animals (Scientific Procedures) Act 1986 Amendment Regulations 2012.

### Preparation of parasites

*S. mansoni* schistosomula were obtained by mechanical transformation of cercariae and cultured as described previously^83^. In brief, snails were washed, transferred to a beaker with water (~50-100 ml) and exposed under light to induce cercarial shedding for two hours, replacing the water and collecting cercariae every 30 min. Cercarial water collected from the beaker was filtered through a 47µm stainless steel Millipore screen apparatus into sterile 50 ml-Falcon tubes to remove any debris and snail faeces. The cercariae were concentrated by centrifugation (800 g for 15 min), washed three times in 1X PBS supplemented with 2% PSF (200 U/ml penicillin, 200 µg/ml streptomycin, 500 ng/ml amphotericin B), and three times in ‘schistosomula wash medium’ (DMEM supplemented with 2% PSF and 10 mM HEPES (4-(2-hydroxyethyl)-1-piperazineethanesulfonic acid)). The cercarial tails were sheared off by ~20 passes back and forth through a 22-G emulsifying needle, schistosomula bodies were separated from the sheared tails by Percoll gradient centrifugation, washed three times in schistosomula wash medium and cultured at 37°C in modified Basch’s medium under 5% CO_2_ in air^83^.

### Single-cell tissue dissociation

Two days after transformation the schistosomula cultured in modified Basch’s media at 37°C and 5% CO_2_ were collected and processed in two separate batches (batch1 and batch2). Schistosomula collected from two different snail batches were considered biological replicates. Data collected as batch3 are ‘technical’ replicates of batch2 given they were collected on the same day and from the same pool of parasites. In each experiment, approximately 5,000 larvae were pooled in 15ml tubes and digested for 30 min in an Innova 4,430 incubator with agitation at 300 rpm at 37°C, using a digestion solution of 750µg/ml Liberase DL (Roche 05466202001). The resulting suspension was passed through 70µm and 40µm cells strainers (Falcon). Dissociated cells were spun at 300 rpm for 5 mins and resuspended in 1X cold PBS supplemented with 20% heat inactivated fetal bovine serum (twice). The resulting cell suspension was co-stained with 0.5µg/ml of Fluorescein Diacetate (FDA; Sigma F7378) to label live cells, and 1µg/ml of Propidium Iodide (PI; Sigma P4864) to label dead/dying cells, and sorted into eppendorf tubes using the BD *Influx*™ cell sorter by enriching for FDA+/ PI- cells^84^. It took 2-3 hours from the enzymatic digestion to generating single-cell suspensions ready for library preparation on the 10X Genomics Chromium platform.

### 10X Genomics library preparation and sequencing

The concentration of single cell suspensions was approximately 500 cells/µl as estimated by flow cytometry-based counting. Cells were loaded according to standard protocol of the Chromium single-cell 3’ kit in order to capture approximately 7,000 cells per reaction (V2 chemistry). However, after sequencing and preliminary analysis, we found the actual number of captured cells was closer to ~1200 cells per experiment. Single-cell libraries were sequenced on an Illumina Hiseq4000 (paired-end reads 75bp), using one sequencing lane per sample. All raw sequence data is deposited in the ENA under the project accession ERP116919.

### Protein-coding genes

*S. mansoni* gene annotation is based on the latest genome assembly (v7, unpublished). The identifier for all genes contains the Smp_ prefix followed by a unique 6-digit number; entirely new gene models have the first digit ‘3’, eg. Smp_3xxxxx. To assign a gene name and functional annotation (used in Tables S4-S6) to ‘Smp_’ identifiers, protein-coding transcript sequences were blasted against SwissProt3 to predict product information (blastp v2.7.0). Some genes also maintained previous functional annotation from GeneDB. Genes lacking predicted product information were named hypothetical genes.

### Mapping and quantification of single-cell RNA-seq

Single-cell RNA-seq data were mapped to the *S. mansoni* reference genome version 7 (https://parasite.wormbase.org/Schistosoma_mansoni_prjea36577) using the 10X Genomics analysis pipeline Cell Ranger (v 2.1.0). The default cut-off provided by Cell ranger was used to detect empty droplets. Approximately 55% of sequenced reads mapped confidently to the transcriptome with an average 297,403 reads per cell. In total 3,513 cells were sequenced, with a median 918 genes expressed per cell.

### Quality control of single-cell data

To filter lower quality cells, the best practices for pre-processing and quality control from the Scater package (version 1.8.4)^85^ were followed. We first created a single cell experiment using *SingleCellExperiment: S4 Classes for Single Cell Data* R package version 1.5.0^86^. Cells that had greater than 30,000 Unique Molecular Identifiers (UMIs) were removed. Although tools are not currently available to determine biological doublets, at the concentrations of cells used in these experiments, the doublet rate is expected to be very low (~1%). In addition, cells with mitochondrial gene expression greater than 3% or cells that expressed fewer than 600 genes per cell were excluded.

We further filtered the data by generating a consensus matrix with the SC3 package (version 1.8.0)^41^ and excluded any clusters with a cluster stability index of less than 0.10. This was done on the basis that cells with low stability index included cells that could not be assigned confidently to a specific cell population. We also excluded clusters containing less than 3 cells due to the limitations of SC3 to capture rare cell types^41^. In total 2,144 cells out of 3,513 cells passed QC. Further exclusion of one ambiguous cluster resulted in a total of 1,918 cells.

### Data normalisation

Data was first clustered with the quickCluster function from scran (version 1.8.4)^87^. The quickCluster function groups cells according to their expression profiles. Cell biases were normalised using the computeSumFactors function. The computeSumFactors function works on the assumption that most genes are not differentially expressed between cells. As such, any differences in expression across the majority of the genes are the result of technical biases in the single-cell dataset and need to be removed^87^. Finally, the normalised expression values were calculated using the normalise function from the scater package (version 1.8.4)^85^.

### Clustering and QC using SC3

The SC3 package (version 1.8.0) was used to cluster and exclude low quality cells from the dataset^41^. For the consensus clustering, SC3 uses the consensus-based similarity partitioning algorithm (CSPA). SC3 constructs a binary similarity matrix using cell labels. When two cells belong to the same cluster, the assigned value is 1; otherwise the value is 0. A consensus matrix is the result of the averaging of all similarity matrices of individual clustering. Based on the consensus matrix, the cells were then clustered using hierarchical clustering using *k* levels of hierarchy where *k* was specified. In the first instance we used a range of values close to the *k* value estimated using the sc3_estimate_k function (*k*=26) from the SC3 package. The stability and quality of the clusters was assessed by visually inspecting the data obtained for the specified *k* value ranges. Clusters with stability index less than 0.10 and/ or less than 3 cells were excluded from further analysis. We continued to re-cluster cells until all clusters had stability values greater than 0.6 and contained more than 5 cells. We also sub-clustered populations of cells that were contained within the same *k* level of hierarchy but appeared to be distinct subpopulations of cells.

### Clustering using Seurat after QC steps

The Seurat package (version 2.3.4) (https://satijalab.org/seurat/) was used to analyse the raw values of QC matrix^42^. First, we normalised using the NormalizeData function from Seurat (http://satijalab.org/seurat/). Following normalisation, we identified highly variable genes using the Seurat FindVariableGenes function using the cut-offs stated in the website: *z*=0.5 and mean expression in the range 0.0125 to 3. We identified 12 clusters (including the ambiguous cluster) using the FindClusters function from Seurat with a resolution of 0.6.

### Identifying marker genes and cluster annotation

To annotate each cluster, we manually inspected the top markers for each of the populations and compared to the top markers curated from the literature (Table S6). We used the top markers identified by SC3 and Seurat packages. SC3 identifies marker genes for each cluster by constructing a binary classifier based on the mean expression values for each gene. The area under the operating characteristic (ROC) curve is used to quantify the confidence for that specific marker. A Wilcoxon signed-rank test is used to assign a P-value to each gene. We relied on high quality marker genes with area under the curve (AUROC) > 0.8, *P* < 10^−5^ and spatial information of those genes to determine the identity of a specific population. We also used the Seurat package to identify marker genes for each population using the function FindAllMarkers, using the likelihood ratio as specified in the Seurat best practices (https://satijalab.org/seurat/).

### Gene Ontology (GO) analysis

The Gene Ontology (GO) annotation *for S. mansoni* was obtained using InterProScan v5.25-64.0 (https://www.ebi.ac.uk/interpro/). GO term enrichment was performed using the weight01 method provided in topGO^88^ v2.34.0 (available at http://bioconductor.org/packages/release/bioc/html/topGO.html) for all three categories (BP, MF, and CC). For each category, the analysis was restricted to terms with a node size of >= 5. Fisher’s exact test was applied to assess the significance of overrepresented terms compared with all expressed genes. The threshold was set as FDR < 0.01.

### STRINGdb Analysis

We used STRINGdb^89^ to identify possible gene interactions that would enable us to differentiate between tegumental clusters. Briefly, the *S. mansoni* V7 gene identifiers for the tegument 2 cluster with AUROC ≥0.7 in Seurat were converted to *S. mansoni* V5 gene identifiers. The V5 gene identifiers were analysed in STRINGdb v11.0^89^. Human, *Caenorhabditis elegans* and *Drosophila melanogaster* orthologs of these genes were identified from WormBase ParaSite^90^.

### Random Forest (RF)

A single-cell dataset published for *Schmidtea mediterranea* comprising 21,610 cells generated using a droplet-based platform^31^ was employed for this analysis. The relevant files were downloaded from https://shiny.mdc-berlin.de/psca/ including the *Schmidtea mediterranea* single-cell data comprising 21,610 cells. The Seurat package (version 2.3.4) was used for all analysis of the *Schmidtea* dataset (https://satijalab.org/seurat/). We only kept cells that expressed at least 200 genes, in a minimum of 3 cells. After QC, 21,612 cells and 28,030 transcripts remained. We normalised using NormalizeData function from the Seurat (http://satijalab.org/seurat/). Following normalisation, we identified highly variable genes using the Seurat FindVariableGenes function using the cut-offs stated in the website: *z*=0.5 and mean expression in the range 0.0125 to 3.

We identified 22 clusters using the FindClusters function from Seurat with a resolution of 0.6. We chose this resolution to capture most of the clusters with biological variability whilst avoiding overclustering. To annotate each cluster, we used the annotation provided by Plass *et al*, 2018^31^.

### Evaluating the Random Forest on the *Schmidtea mediterranea* dataset

We accessed the transcriptome reference (version 6) for the asexual strain of *Schmidtea mediterranea* from planmine^91^. This version is a Trinity *de novo* transcript assembly^92^. We used orthoMCL^93^ to find 1:1 ortholog genes between *S. mediterranea* and *S. mansoni* by: (i) collapsing Smp and dd_Smed genes to their root names and choosing clusters with a single *Schmidtea* and *Schistosoma* gene; and (ii) removing haplotype Smp genes where doing so would reduce a multiple Smp set to a single Smp; (iii) If a single Smp (after all the above checks) contained multiple *Schmidtea* genes, we randomly selected one of the *Schmidtea* genes only if it did not map to more than one orthologue cluster. This gave us a set of *Schmidtea-Schistosoma* orthologous gene-pairs. All *Schistosoma* genes were then replaced in the *Schistosoma* single-cell matrix with their *S. mediterranea* orthologs.

We first evaluated the RF classifier on the *Schmidtea* dataset. We used R package randomForest (version 4.6-14) to train the training set using 500 trees. The RF is a supervised learning method that builds decision trees, trained with a defined set of features (genes). The training set was built using cells from the 22 clusters in the *Schmidtea* dataset with a maximum of 70% of cells per cluster. As a first RF test, the training set (70% of cells per cluster) was used to assign a cluster label for the test set (remaining 30%) of the same dataset. We assigned a class to each cell when a minimum of 16% of trees in the forest converged onto a decision.

To then use the RF classifier on the *Schistosoma* data set, a training set was built using cells from the 22 clusters in the *Schmidtea* dataset with a maximum of 70% of cells per cluster. This training set was used to assign labels to the *Schistosoma mansoni* cells using the RF package^94^. The RF decision trees were trained with a defined set of common 692 orthologous genes between *S. mansoni* and *S. mediterranea*.

### Conserved Schmidtea-Schistosoma orthologous markers

To identify conserved *Schmidtea-Schistosoma* one-to-one orthologs, we first identified a high confidence set of one-to-one orthologs. For each *S. mansoni* predicted protein, we identified the *S. mediterranea* BLASTP^95^ hits, and similarly identified the *S. mansoni* BLASTP hits for each *S. mediterranea* protein. If a *S. mansoni* gene had haplotypic copies in the *S. mansoni* V7 assembly, we only considered the *S. mansoni* copy on an assembled chromosome, and discarded the allelic copies of the gene from haplotypic contigs. We considered *S. mansoni* and *S. mediterranea* genes to be one-to-one orthologs if they were each other’s top BLASTP hits, with BLAST E<0.05, and the BLAST E-value of the top BLASTP hit was 1e+5 times lower than the BLAST E-value for the next best hit. This gave us 4764 one-to-one *S. mansoni-S. mediterranea* orthologs. These orthologs were used to find conserved orthologous markers.

To identify conserved orthologous markers, we filtered the 4764 1:1 orthologues to retain only those for which both the *Schistosoma* and *Schmidtea* genes were identified as Seurat markers with Seurat P-value ≤1e-30, using the Seurat *Schmidtea* clusters from Plass *et al* 2018^31^. If a *Schistosoma/Schmidtea* gene was in more than one cluster, we only considered the cluster for which it had the lowest (most significant) Seurat P-value.

### *In situ* hybridization (ISH)

Fluorescence *in situ* hybridization (FISH) and whole-mount colorimetric *in situ* hybridization (WISH) were performed following previously established protocols^14,15,49^ with modifications specific to schistosomula. Schistosomula were killed with ice-cold 1% HCl for 30–60 s before fixation. Schistosomula were fixed for ~0.5-1 hour at room temperature in 4% formaldehyde, 0.2% Triton X-100%, 1% NP-40 in PBS. Adult parasites were fixed for 4 hours in 4% formaldehyde in PBSTx at room temperature. After fixation, schistosomula and adults were dehydrated in methanol and kept in −20°C until usage. Parasites were rehydrated, permeabilised by 10 µg/mL proteinase K for 10-20 min for schistosomula or 20 µg/mL proteinase K for 30 min for adults, and fixed for 10 mins immediately following proteinase K treatment.

For hybridization, DIG-riboprobes were used for single FISH and WISH, and FITC-riboprobes were used for double FISH. Anti-DIG-POD and anti-FITC-POD antibodies were used for FISH at 1:500-1:1000, and anti-DIG-AP antibody was used for WISH. Anti-DIG-POD and anti-DIG-AP antibodies were incubated overnight at 4°C and anti-FITC-POD was incubated for ~4 hours at room temperature before overnight incubation at 4°C. For FISH, 2-3 independent experiments were performed, and ~5-10 worms were analysed for each experiment. For adult FISH and WISH, two independent experiments were performed, with each experiment containing ~5 male and ~5 females. Primers used for cloning a fragment of marker genes and riboprobe generation are listed in Table S7.

### Immunostaining and labeling

Anti-acetylated ɑ-tubulin antibody (6-11B-1, Santa Cruz) was incubated at 1:500 in blocking solution (5% Horse serum, 0.5% Roche Western Blocking Reagent in TNTx). Secondary antibody (anti-mouse Alexa Fluor 633, Invitrogen) was used at 1:250-1:500 and was incubated overnight at 4°C. For lectin labeling, fluorescein succinylated wheat germ agglutinin (sWGA) (Vector Labs) was used at 1:500 dilution in a blocking solution overnight at 4°C. Fluorescent dextran was used to label tegument cells^6^. Briefly, schistosomula were transferred to 20µm mesh in order to flush out as much media while retaining parasites inside the mesh. 2.5mg/ml dextran biotin-TAMRA-dextran (ThermoFisher Scientific, D3312) was added to the mesh and parasites transferred into a 1.7ml tube. Immediately after the transfer, schistosomula were vortexed for ~2-4 minutes at 70% vortex power, transferred back to 20µm mesh and flushed with schistosomula fixative (4% formaldehyde, 0.2% Triton X-100%, 1% NP-40 in PBS) before fixing.

### Imaging and image processing

Schistosomula FISH images were taken using an Andor Spinning Disk WDb system (Andor Technology). Adult FISH images were taken using a Zeiss LSM 880 with Airyscan (Carl Zeiss) confocal microscope. Colorimetric WISH images were taken using AxioZoom.V16 (Carl Zeiss). Imaris 9.2 (Bitplane) and Photoshop (Adobe Systems) was used to process acquired images of maximum intensity projections (of z-stacks) and single confocal sections for linear adjustment of brightness and contrast.

### Calculating cell numbers in schistosomula

Cercariae and parasites at 0, 24 and 48 hr post-transformation were fixed in 5% (v/v) formaldehyde 4% (w/v) sucrose in PBS for 15 min (throughout staining worms were in 1.5 ml microfuge tubes and spun down 2 min 500G when exchanging solutions). The parasites were then permeabilised in 10% (w/v) sucrose, 0.5 % Triton-X 100 (v/v) for 10 min. Parasites were either stored at 4°C in 2% formaldehyde in PBS, or stained immediately. Staining was in low light level conditions to minimise photobleaching. 1 µg/ml DAPI in PBS was added for 10 min, then parasites were post-fixed in 10% formaldehdye in PBS for 2 min, washed in 1X PBS, then resuspended in 0.4X PBS in ddH_2_O (to discourage salt crystals). 10 µl parasites were pipetted onto a glass slide and excess liquid drawn away with whatman filter paper. 10 µl ProLong Gold antifade mountant was added to the sample and a glass coverslip dropped over gently. Slides were left at room temperature overnight to set before imaging. A Zeiss LSM 510 Meta confocal microscope was used in conjunction with the Zen software to take a series of Z stacks, imaging 3 individual worms from each timepoint.

Z stack images were imported into ImageJ software (Import>image sequence) then converted to RGB and split by color (Color>split channels) and the blue channel used for further processing. Using the metadata associated with the file the scale properties were adjusted. The image was cropped if necessary to show only one parasite. The threshold was set to remove any background. The signal above threshold was measured for the whole image stack (image can be inverted and converted to 8 bit for this purpose). The ROI manager was used to measure individual cell nuclei throughout the Z stack by drawing around the cell on each image of the stack where present. This was imported to the threshold filtered stack and the area measured. 10 nuclei that were clearly defined and of diverse location and size were measured for each worm to obtain an average nuclei size and signal. In all cases, Z was used as well as X and Y to account for the full volume of the nuclei. The total volume for above threshold signal in the worm was divided by the average nuclei size to obtain an estimate for cell number.

## Acknowledgements

The work at WSI was supported by Wellcome (award numbers 206194 and 107475/Z/15/Z). PAN is an investigator of the Howard Hughes Medical Institute. *B. glabrata* snails used in the United States were provided by the NIAID Schistosomiasis Resource Center of the Biomedical Research Institute (Rockville, MD) through NIH-NIAID Contract HHSN272201700014I for distribution through BEI Resources. We thank the Sequencing and Informatics core facilities at WSI for their contribution. We also thank the following: Gal Horesh for initial technical assistance optimising the dissociation conditions; Catherine McCarthy and Simon Clare for technical support with animal infections and maintenance of the *S. mansoni* life cycle; David Goulding and Claire Cormie at the Electron and Advanced Light Microscopy facility; Jennie Graham and Sam Thompson at the Cytometry Core Facility; Nancy Holroyd, Mandy Sanders, Elizabeth Cook and Nathalie Smerdon for facilitating the submission of 10X samples; and Matthew Jones for 10X training and library preparations. The authors thank Dr. Shristi Pandey for sharing the random forest code used in this work and Dr. Mireya Plass for sharing the planaria dataset.

---

**Supplementary Figure 1.**
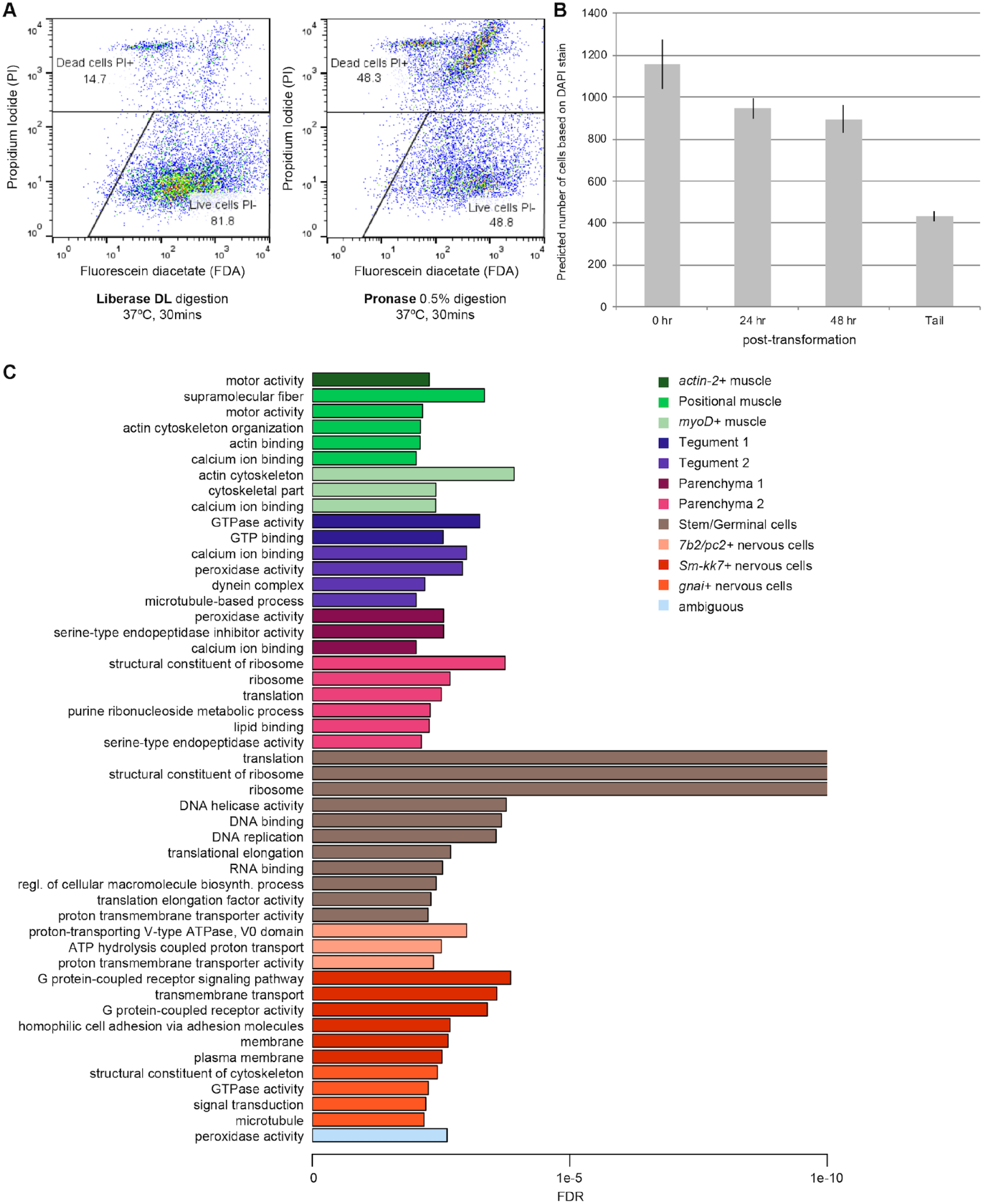
(A) Comparison between protocols to dissociate schistosomula. Flow cytometry-based assessment of dissociation with either Liberase DL (750μ1/ml) (left) or Pronase 0.5% (right) revealed that the former led to more live cells than the latter. (B) Predicted number of cells that comprises an *in vitro-transformed* schistosomulum. The bar chart shows the number of cells counted in schistosomula immediately after mechanical transformation (0 hr, i.e. cercaria head), after one day (24 hr) and two days (48 hrs) in culture. ‘Tail’ represents the number of cells counted in the tail detached from the cercaria during the mechanical transformation. Mean of number of cells counted in 3 schistosomula per timepoint (C) Significantly enriched GO terms for the marker genes in each cell cluster. Plot showing terms with FDR< 0.01 from a Fisher’s exact test and coloured by cell types. The x-axis indicates -log! 0FDR values, where an arbitrary maximum value of IO was set.

**Supplementary Figure 2.**
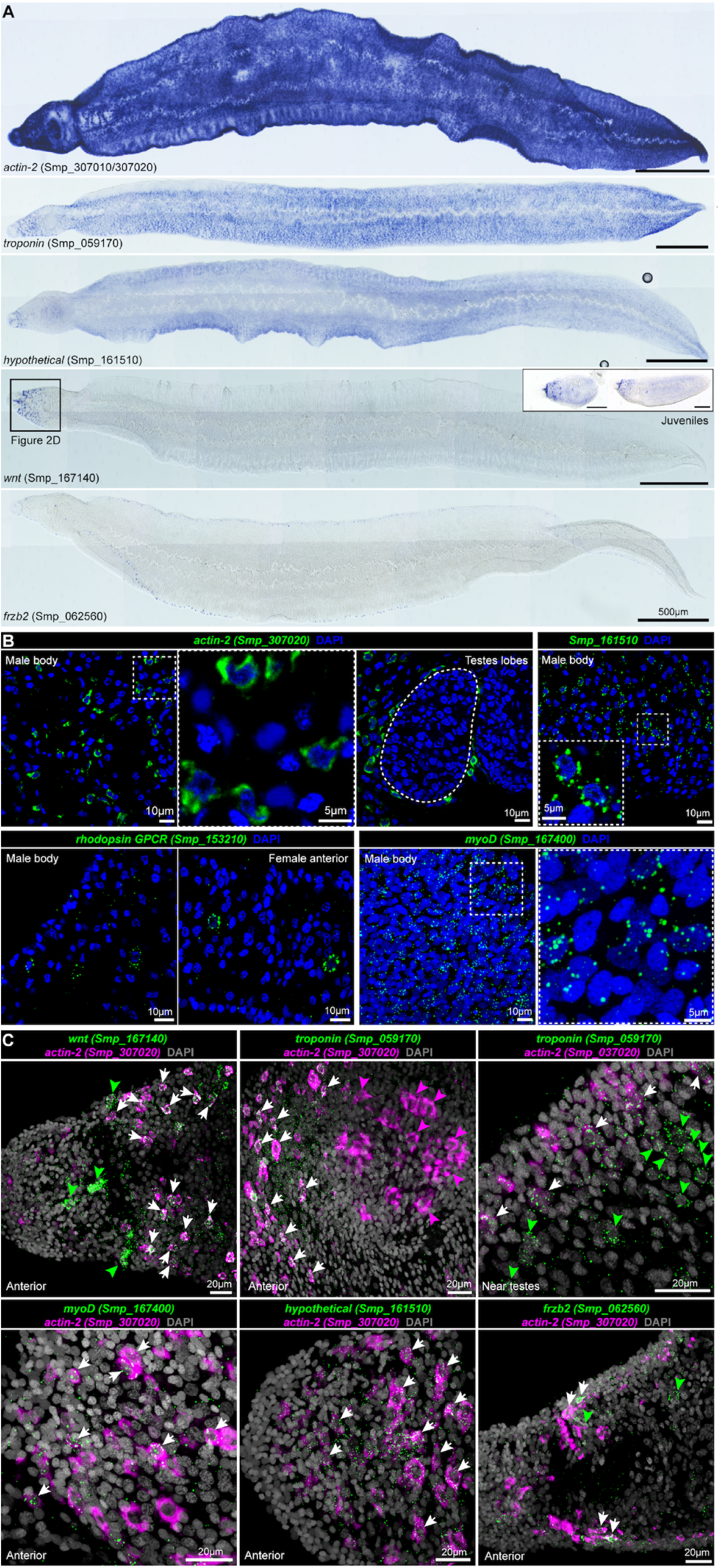
(A) WISH of indicated markers and signalling molecules enriched in a subset of muscle cells in adult schistosomes. For *wnt*, the boxed region is shown in Figure 2D. On the right, WISH experiment shows that *wnt* expression is conserved in the anterior end of juvenile parasites collected from mice 3 weeks post-infection. (B) FISH of muscle markers in indicated regions of the adult worms. (C) Double FISH of selected muscle markers. White arrows: double positive cells; green arrowheads: single positive cells expressing genes indicated in green; magenta arrowhead: single positive cells expressing genes indicated in magenta. Magnified single confocal sections are shown for the dotted box area to the right of the image.

**Supplementary Figure 3.**
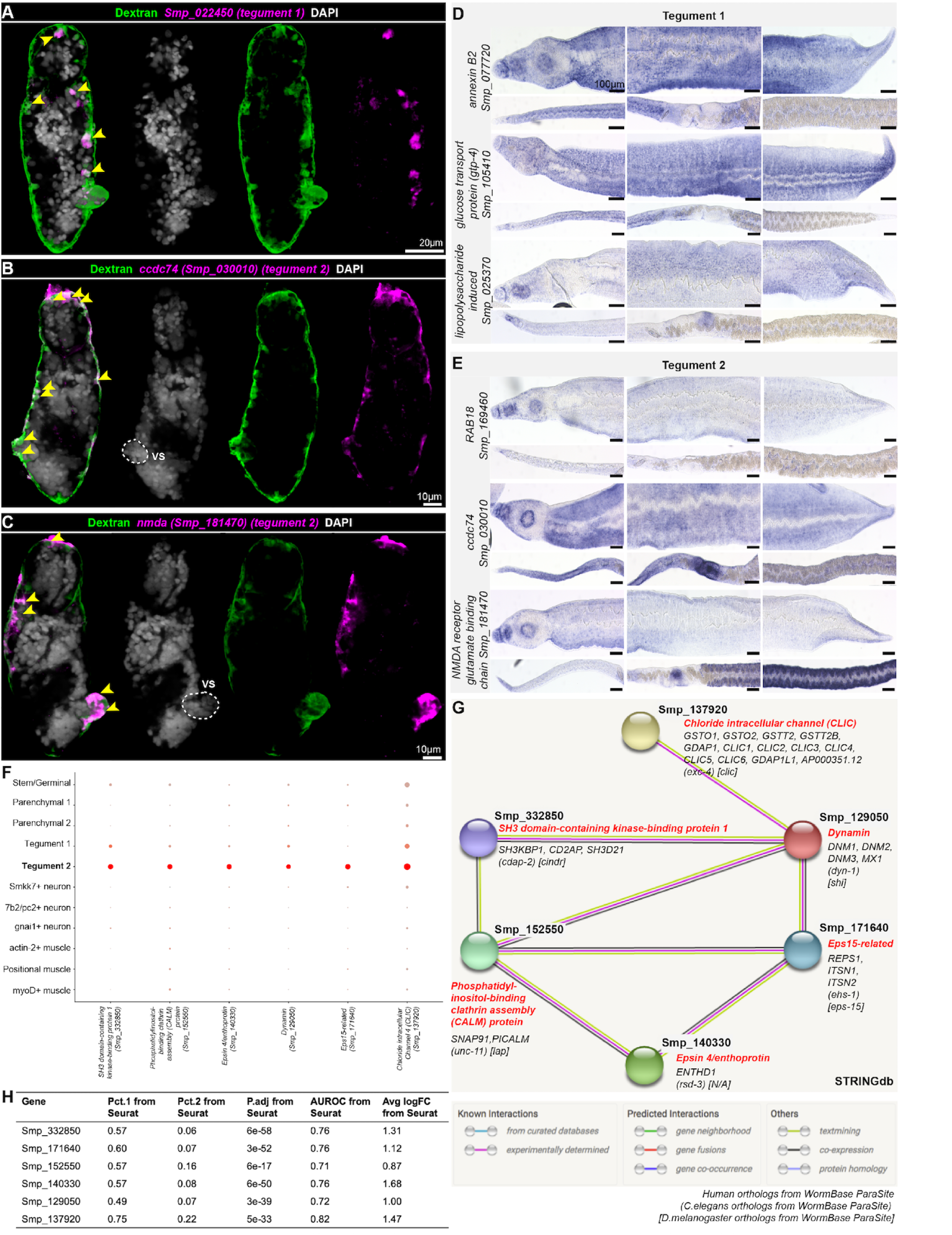
(A-C) Dextran labelling in schistosomula shows co-localisation with Tegument 1 and 2 markers. Yellow arrowheads indicate cells positive for both dextran and tegument marker. VS: ventral sucker. (D-E) WISH of Tegument I and Tegument 2 maker genes in adult parasites. Scale bar: I 00 μm (F) Expression profile of tegument 2 marker genes used for STRINGdb analysis. (G-H) Prediction of biological processes enriched in Tegument 2 relative to Tegument 1. We identified genes that are strong markers (with AUROC 2:0. 7 in Seurat) for Tegument 2 but not for Tegument 1. The set of genes shown is the largest connected component, i.e. set of predicted interacting genes in the STRINGdb results. The interaction cluster included several genes related to clathrin-mediated (receptor-mediated) endocytosis. These included phosphatidylinositol-binding clathrin assembly protein (CALM), Eps 15-related, and epsin-related genes.

**Supplementary Figure 4.**
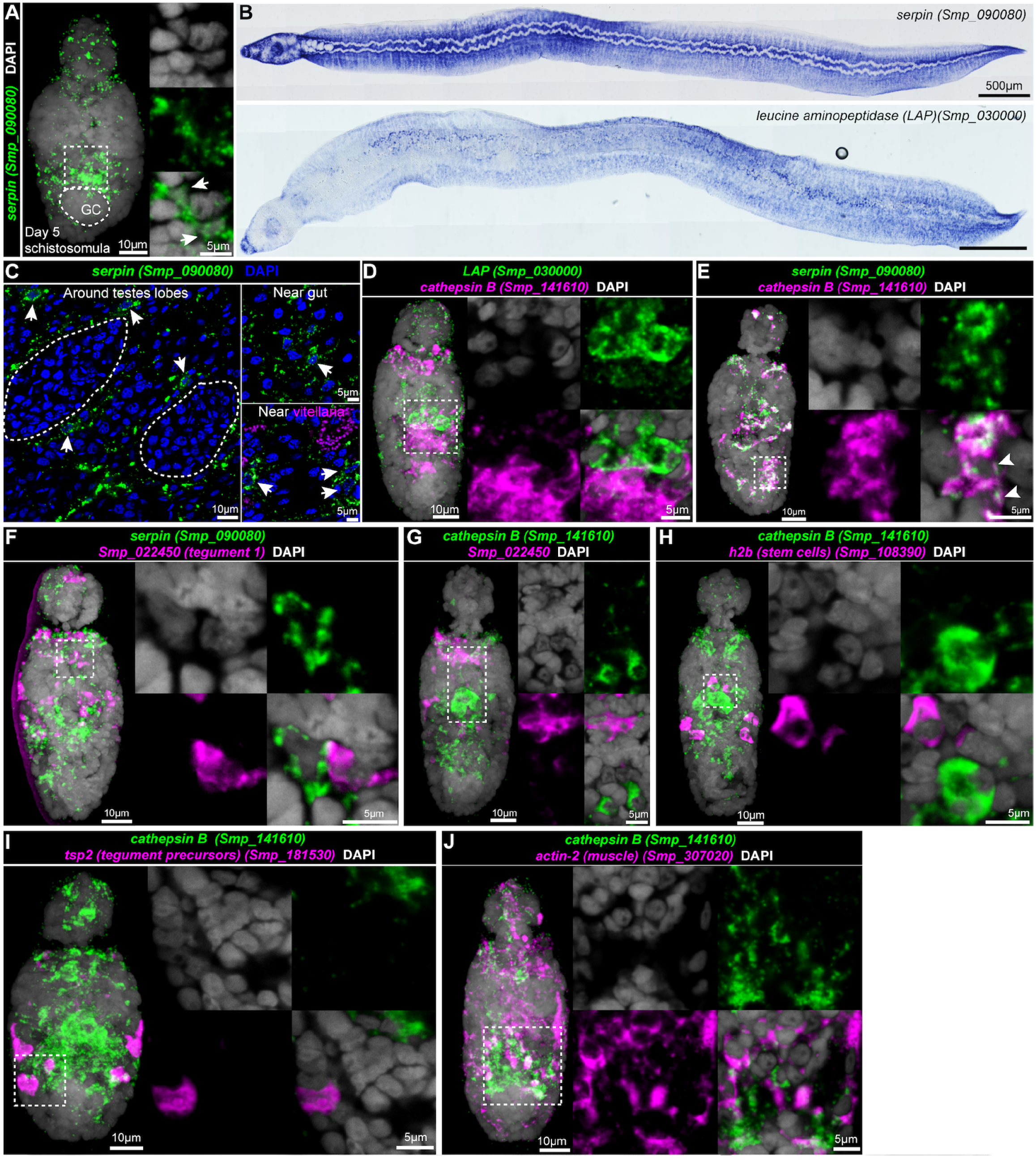
(A) *serpin* FISH in five-day old schistosomula. MIP of whole worm is shown on the left, and single magnified confocal section from the dotted box is shown on the right. White arrows indicate a positive cell that has long cytoplasmic processes. (B) WISH of *serpin* and *lap* in adult parasites; *lap* is expressed in the worm parenchyma as well as in the gut. (C) Single confocal sections showing FISH of *serpin* in different regions of the worm. White arrows indicate single positive cells. (D-J) Double FISH of parenchymal cell markers and other indicated cell type markers in two-day old schistosomula. Parenchymal cell markers do not co-localise with the tegument cells (F-G), stem cells (H), or tegument precursors (I) but show some co-localisation with muscle cells (J). MIP is shown for the whole worm on the left, and single confocal sections from the dotted box are shown on the right.

**Supplementary Figure 5.**
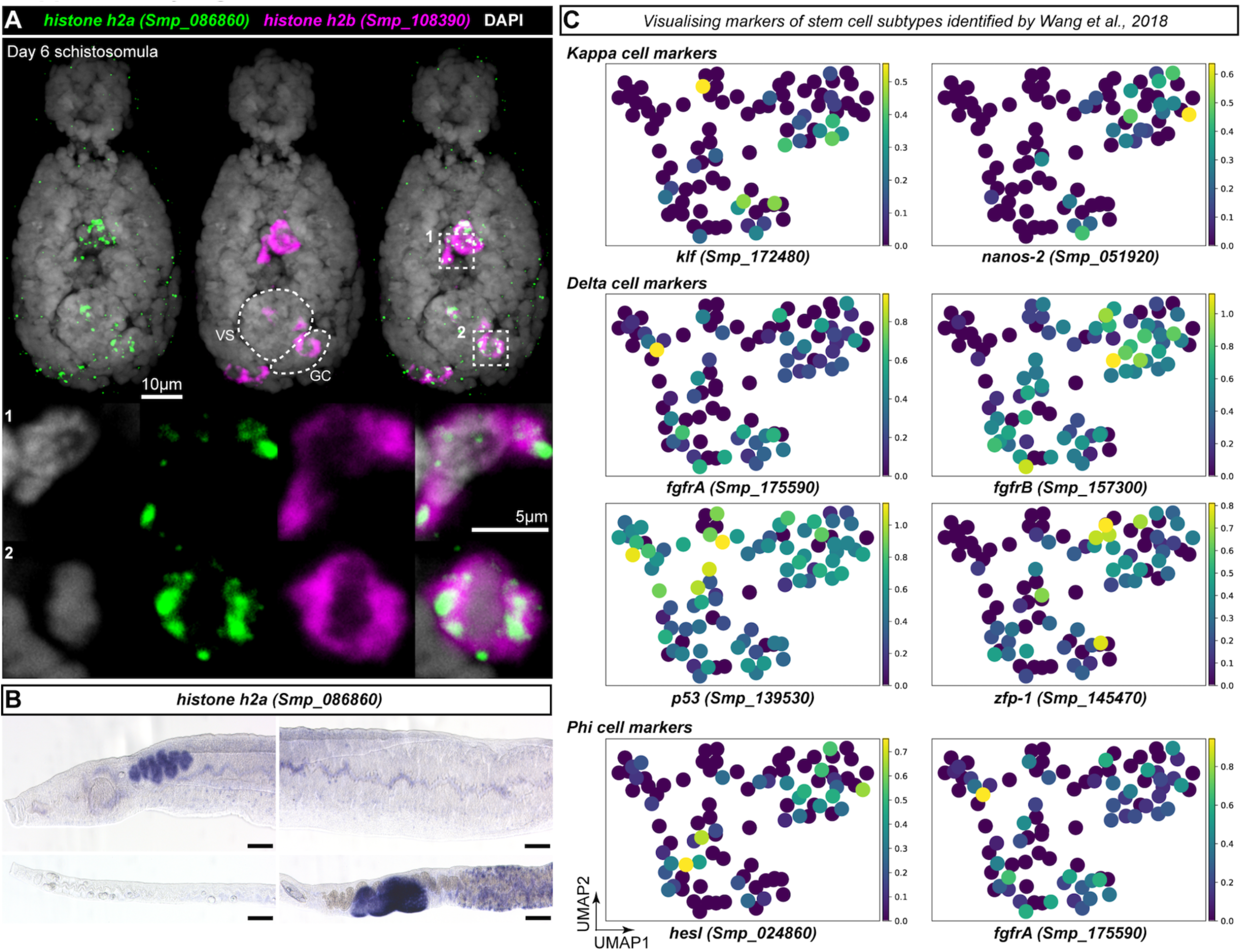
(A) Double FISH of *h2a* identified in our dataset, and *h2b*, a known validated schistosome stem cell marker in six-day old schistosomula. *h2a+* cells co-express *h2b* in both the soma as well as in the germinal cell cluster. Top: MIP for whole worm; Bottom: single confocal magnified sections from the dotted box regions. (B) WISH of *h2a* in adult parasites show expression in gonads and somatic cells, consistent with calmodulin and other previously characterised stem cell genes. (C) UMAP plots showing the expression of marker genes for three stem cell populations identified by Wang et al., 2018. Color scale of the UMAP plots ranges from yellow (high expression) to dark blue (no expression).

**Supplementary Figure 6.**
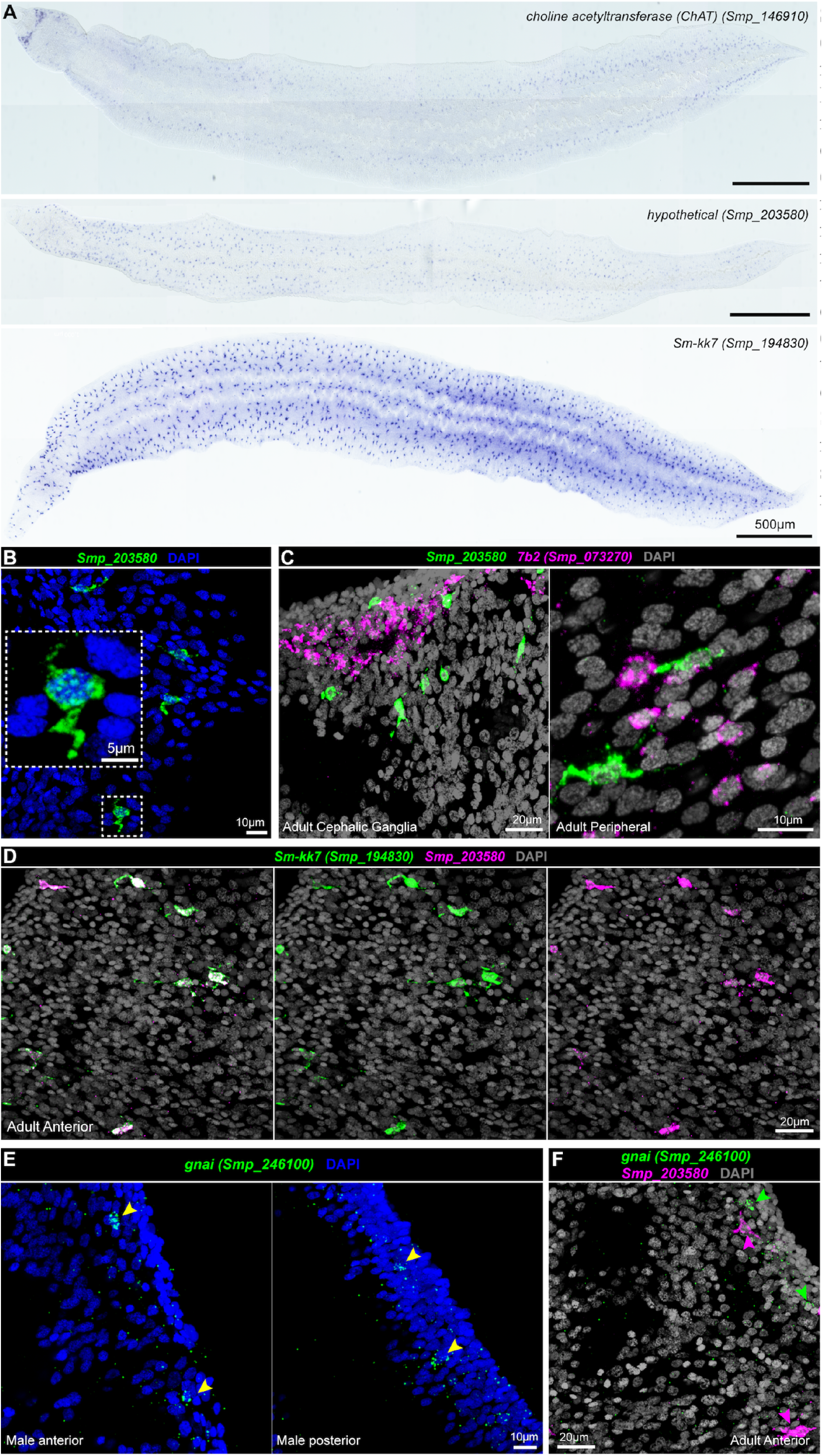
(A) WISH of indicated neuronal markers in adult parasites. (B) FISH of Smp 203580 in adult male soma (mid-body) shows long cellular processes in each cell. (C-D) Double FISH in adults reveals that (C) Smp 203580 does not co-localise with pan-neuronal marker *7b2*, but (D) nearly all cells that express Smp_203580 co-express *Sm-kk7* (white signal). (E-F) *gnai* is expressed throughout the body of the adult worm (E), but does not co-localise with Smp_203580 (F). Green and magenta arrowheads indicate single positive cells for respectively labelled genes.

